# Population structure and genetic variance among local populations of an non-native earthworm species in Minnesota, USA

**DOI:** 10.1101/2022.10.11.511724

**Authors:** Bastian Heimburger, Andreas Klein, Alex Roth, Stefan Scheu, Nico Eisenhauer, Ina Schaefer

**Affiliations:** J.F. Blumenbach Institute of Zoology and Anthropology, University of Göttingen, Untere Karspüle 2, 37073 Göttingen, Germany; Department of Forest Resources, University of Minnesota 115 Green Hall, 1530 Cleveland Ave. N., St. Paul, MN 55108; Friends of the Mississippi River, 101 E 5th St., Suite 2000, St. Paul, MN 55101; German Centre for Integrative Biodiversity Research (iDiv), Puschstr. 4, 04103 Leipzig, Germany; Institute of Biology, Leipzig University, Puschstr. 4, 04109 Leipzig, Germany; Centre of Biodiversity and Sustainable Land Use, Büsgenweg 1, 37077 Göttingen, Germany

**Keywords:** population genetics, invasion, earthworms, multiple introductions, diffusion dispersal, *L. terrestris*

## Abstract

A variety of human activities have been identified as driving factors for the release and spread of invasive earthworm species in North America. Population genetic markers can help to identify locally relevant anthropogenic vectors and provide insights into the processes of population dispersal and establishment. We sampled the invasive European earthworm species *Lumbricus terrestris* at nine sites and several bait shops within the metropolitan area of Minneapolis-St. Paul in Minnesota, USA. We used microsatellite markers to infer genetic diversity and population structure, and 16S rDNA to address multiple introduction events, including bait dumping, which is a common source of *L. terrestris* introductions into the wild. Our results indicate multiple introductions but not from current bait dumping. Overall, genetic structure was low and earthworms >5000 m apart were genetically differentiated, except for one sampling location, indicating jump-dispersal followed by population establishment. Further, earthworms at one location north of Minneapolis established from one or few founder individuals, suggesting that earthworm invasions are ongoing. We therefore encourage further monitoring of earthworm populations using molecular markers, in order to disentangle the different human-related vectors contributing to the spread of earthworms and their establishment, which is essential to develop adequate management strategies.

## Introduction

The invasion of North American forests by European earthworms is indicated by short- and long-term environmental changes they cause. These lumbricid earthworms actively reduce the litter layer on the soil surface and mix the litter and humus layers, which indirectly reduces soil carbon and nitrogen over the long-term and depletes other soil minerals (Crumsey et al. 2015; Resner et al. 2015; Ferlian et al. 2020). Through their burrowing activity, they also influence the availability of nutrients to plants by disrupting mycorrhizal networks (Paudel et al. 2016; Dobson et al. 2017). Additionally, their presence in North American forests has facilitated the invasion of non-native plant species by selective seed predation and changes in nutrient availability (Nuzzo et al. 2009; Drouin et al. 2014; Nuzzo et al. 2015; Craven et al. 2017). In consequence, non-native earthworms indirectly and directly affect nutrient availability to soil and aboveground communities and have irreversible consequences for forest ecosystems (Eisenhauer et al. 2007; Fisichelli et al. 2013; Fisichelli and Miller 2018; Ferlian et al. 2018; Frelich et al. 2019). Thus, it is important to identify potential introduction pathways in order to protect undisturbed native communities and to prevent future invasions.

The role of human transport for earthworm introductions and dispersal into the wild is well established. Disposal of earthworms used as fishing bait and passive transport by vehicles along roads have been identified as central drivers (Holdsworth et al. 2007; Keller et al. 2007; Cameron et al. 2007, 2008; Klein et al. 2017, 2020). Further, human disturbances seem to provide suitable conditions for non-native earthworm invasions, such that active or historic human settlements, construction sites, agricultural use, and recreational areas and fishing sites contain more non-native earthworms than wilderness and undisturbed sites (Hendrix et al. 2006, 2008; Callaham et al. 2016; Rogers and Collins 2017; Arcese and Rodewald 2019). The identification of vulnerable sites and vectors for earthworm invasions is necessary to formulate containment guidelines, and knowledge of the local earthworm population structure is required to understand how invasive earthworms establish viable populations.

Introduction from distant or locally adapted populations can increase genetic diversity, allowing populations to mitigate founder-effects and increase the potential of adaptation to local environmental conditions (Lawson Handley et al. 2011). Genetic studies of non-native earthworms in northern North America clearly link them to their European or Asian sources (Gailing et al. 2012, Fernandez et al. 2016, Schult et al. 2016) and demonstrate that genetic diversity of populations is generally high (Gailing et al. 2012, Porco et al. 2013, Schult et al. 2016, Keller et al. 2020), particularly in areas used by humans (Cameron et al. 2008, Klein et al. 2017). This indicates multiple introductions but leaves open the question whether these populations are permanent or the result of continuous re-introductions by human transport. Molecular markers are needed to analyse the population structure of free-living earthworms, to determine the genetic variability in populations, to identify hotspots of multiple introductions, and to infer patterns of dispersal or the direction of transport.

In this study, we investigated the population structure of *Lumbricus terrestris* Linnaeus, 1758 within the metropolitan area of Minneapolis-St. Paul, Minnesota, USA. This species has been reported in Minnesota since the second half of the 20th century (Gates 1978, Reynolds et al. 2002), and thus is ideal for studying the genetic structure within and among populations. Our sampling sites included recreational and agricultural sites, and were both distant and close to road networks. We sampled earthworms at three locations within a radius of about 50 km, and three subpopulations at each location. Individual earthworms were genotyped with eight microsatellite markers to investigate the incidences and the range of dispersal, as well as genetic diversity within populations. We also included earthworms from five bait shops within the sampling area in order to assess the role of bait dumping as a potential source of free-living earthworm populations. To address multiple introduction events, we sequenced the mitochondrial 16S rDNA gene of all genotyped individuals. First, we analysed population differentiation of *L. terrestris*, which is a poor disperser that concentrates its activity within a small radius around its burrows (Sims and Gerard 1999; Tiunov et al. 2006; Addison 2009; Potvin and Lilleskov 2017), and therefore relies on passive dispersal to cover larger distances. Second, we analysed population structure to check if alleles are shared among locations, which indicates admixture. Third, we analysed population structure based on 16S rDNA to infer the number of genetic lineages at sampling sites and to identify potential hotspots of multiple introductions.

## Methods

### Collection sites, collecting methods and DNA isolation

Three populations of *L. terrestris* were collected in forest habitats between May and July 2014 at a distance of 25 – 75 km from the urban area of Minneapolis-St.-Paul, MN, USA (Table 1, Fig. 1). At each sampling site (labelled as subpopulation _1), two additional replicates were taken at 5 m (labelled as subpopulation _2) and 5000 m (labelled as subpopulation _3) distance, dividing each population into three subpopulations. Adult and juvenile earthworms were collected from plots of 0.5 × 0.5 m using mustard extraction (Gunn 1992) and hand-sorting. Sampling site MN_0 was about 70 km south of Minneapolis and located in a woodland, intersected by a single gravel road that was surrounded by farmland, near Interstate 35 (Fig. 1). Sampling site MN_50 was located in a suburban area of Minneapolis approximately 100 km north-east from MN_0. Several lakes and parks for recreational activities are located in this area. The third sampling location MN_100 was about 100 km north-west from MN_0 and located in an area characterised by agricultural fields and various lakes that are frequently visited by the local population for fishing and camping, and only accessible by small roads. The sampling site of MN_100_3 was located on a ridge that slopes down to the St. Croix River.

**Figure 1.**
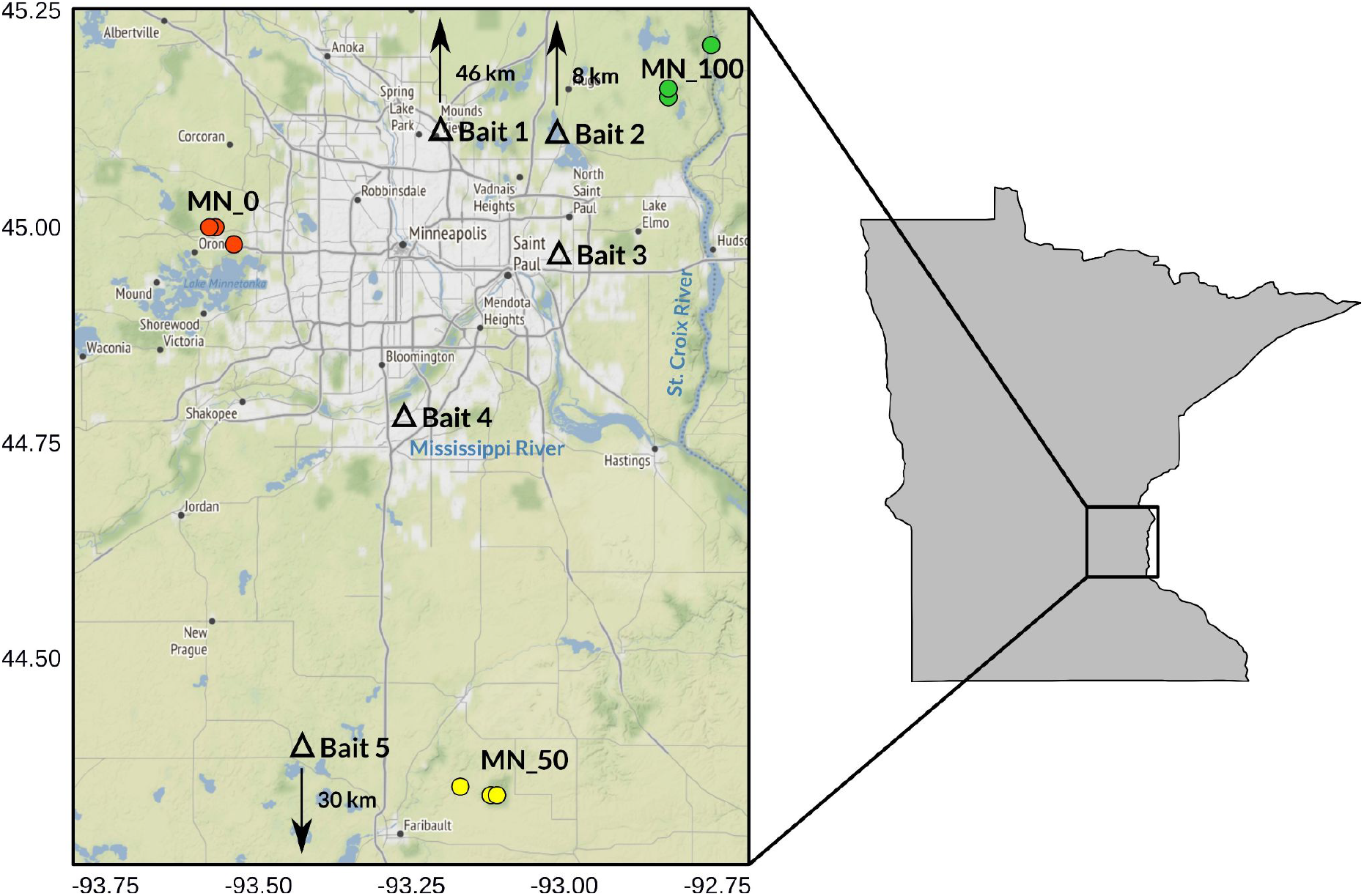
Sampling locations of *Lumbricus terrestris* in Minnesota, USA. Individuals were sampled around Minneapolis-St. Paul (large map) in Minnesota (grey map) in 25 and 75 km distance. Replicates were taken at 5 m and 5 km distance at each sampling site. Triangles indicate locations of bait shops.

To investigate the importance of bait dumping as potential source for introductions, *L. terrestris* fishing baits were included from five different bait shops (five individuals per bait shop) within the sampling region. One centimetre of tail tissue of each sample was shipped from the United States to Germany for barcoding and further molecular analyses. Remaining body parts were stored as vouchers at the University of Minnesota (Department of Forest Resources).

### 16S rDNA sequencing and genotyping

Total genomic DNA was extracted using the Qiagen Blood and Tissue kit (Qiagen; Hilden, Germany) following the manufacturer’s protocol. To verify species identity of juvenile individuals, an 800 bp fragment of 16S rDNA was amplified using the primers 16S-LumbF and Ho_16Sra (Pèrez-Losada et al. 2009), which contain a barcoding region that reliably distinguishes lumbricid species (Bienert et al. 2012). All PCR reactions for sequencing were performed in 25 μl volumes containing 11.75 μl ultrapure H_2_O, 1.25 μl BSA (~4%), 2.5 μl Buffer with KCl, 1 μl dNTPs (10 mM), 3.5 μl MgCl_2_ (25 mM), 0.5 μl Taq polymerase (5 U/μl; Thermo Scientific; Dreieich, Germany), 1 μl of each primer (10 mM), and 2.5 μl template DNA. The PCR protocol consisted of an initial activation step at 95°C for 3 min, 40 amplification cycles (denaturation at 95°C for 30 sec, annealing at 53°C for 60 sec, and elongation at 72°C for 60 sec), and a final elongation step at 72°C for 10 min. PCR products were analysed with multi-capillary electrophoresis using QIAxcel (Qiagen; Hilden, Germany). Some of the PCR products performed inadequately in terms of amplification success and PCRs were repeated in 25 μl volumes containing 2 μl ultrapure H_2_O, 1.5 μl MgCl_2_ (25 mM), 12.5 μl HotStarTaq Mastermix (Genaxxon; Ulm, Germany), 2 μl of each primer (10 mM), and 5 μl template DNA. This PCR protocol consisted of an initial activation step at 92°C for 15 min, 40 amplification cycles as described above but with an annealing temperature of 56°C. PCR products were checked using QIAxcel and positive products were purified using the QIAquick PCR Purification kit (Qiagen; Hilden, Germany) and were either sequenced at the Göttingen Genome Sequencing Laboratory (University of Göttingen) or by Macrogen Europe Inc. (Amsterdam, The Netherlands). All 16S rDNA sequences were checked and ambiguous positions were corrected using electropherograms in Geneious 8.0.5 (https://www.geneious.com), and are available with the accession numbers ON506389 to ON506413 (bait shop individuals) and ON506269 to ON506388 (field collected animals) at NCBI GenBank (www.ncbi.nlm.nih.gov).

We amplified eight highly polymorphic microsatellite loci from *L. terrestris* (LTM 026, LTM 128, LTM 163, LTM 165, LTM 187, LTM 193, LTM 252, and LTM 278; Velavan et al. 2007), using Taq polymerase (Thermo Scientific; Waltham, MA, USA) or the HotStarTaq Mastermix (Genaxxon; Ulm, Germany) following the protocol of Velavan et al. (2007) and Klein et al. (2017). Genotyping of multi locus genotypes (MLGs) was performed at the Department of Animal Sciences, Veterinary Medicine, University of Göttingen, using FAM-labelled forward primers (Sigma Aldrich; St. Louis, MO, USA). Microsatellite profiles were analysed with Geneious 8.0.5 and checked with MICROCHECKER v2.2.3 (http://www.microchecker.hull.ac.uk/; van Oosterhaut et al. 2004) for potential scoring errors.

### Population structure analyses

All 16S rDNA sequences were blasted using the nucleotide BLAST algorithm provided by NCBI GenBank to verify species identity of *L. terrestris*, which was particularly relevant for juvenile individuals included in this dataset. All sequences were corrected by eye and aligned in AliView v1.25 (Larsson 2014) using default settings and remaining wobble positions were conservatively corrected by eye i.e., we manually assigned the nucleotide that was present in all other sequences at this position in the alignment. The alignment was trimmed to the shortest sequence (516 bp) and collapsed to haplotypes with FaBox (Villesen 2007). Population structure was assessed with a haplotype network based on observed p-distances in R (R Core Team, 2020) using the ape and pegas packages (Paradis 2010; Paradis and Schliep 2019). Additionally, we calculated a Maximum Likelihood tree with 1,000 bootstrap replicates to check phylogenetic relatedness of 16S haplotypes using the ape and phangorn packages (Schliep 2011; Paradis and Schliep 2019).

Population genetic analyses of microsatellites were performed using the adegenet package (Jombart 2008, Jombart and Ahmed 2001) in R. All earthworms from bait shops were assigned to a single population (Bait_Shop, 25 individuals) due to their high genetic similarity. For each microsatellite marker, we calculated the mean number of alleles (Na), and observed (Ho) and expected (He) heterozygosity. We tested for significant deviations from expected heterozygosity using the Bartlett test of homogeneity and for significant deviations from Hardy-Weinberg equilibrium (HWE) using a chi-square test. To estimate population differentiation and inbreeding, we calculated the standard diversity indices F_ST_ and F_IS_ (fixation and inbreeding index, Weir and Cockerham F statistics), and pairwise F_ST_ values for four populations (MN_0, MN_50, MN_100 and Bait_Shop) and ten subpopulations (three subpopulations per population and Bait_Shop) using the hierfstat package (Goudet and Jombart 2020). To identify at which hierarchical level the most genetic variation existed, we performed an Analysis of Molecular Variance (AMOVA) using the poppr package v2.9.3 (Kamvar et al. 2014, 2015), and a Monte-Carlo test with 9,999 replications to test for significance. To identify genetic clusters, we did a Discriminant Analysis of Principal Components (DAPC, Jombart et al. 2010), using the xvalDAPC function for cross validation to decide how many axes to retain. This approach partitions genetic variation into a between- and a within-group component, and maximises the first component, thereby reducing the within-group variability, without relying on population genetic models.

We investigated population genetic structure using STRUCTURE v2.3.4 (Pritchard et al. 2000) which, in contrast to DAPC, assigns population to groups that best represent Hardy-Weinberg equilibrium, using a Bayesian sampling approach. Several preliminary runs were performed using various model configurations for optimal prior settings using the LOCPRIOR option. All other settings remained in default mode. Each run comprised ten iterations of the value of K that ranged between 1 and 10 with 10^5^ MCMC burn-in steps and 10^6^ replicates. The most likely number of genetic clusters (ΔK) for all runs of K was calculated in STRUCTURE HARVESTER (Evanno et al. 2005). The greedy algorithm in CLUMPP v1.12 (Jakobsson and Rosenberg 2007) was used to obtain a single mode of K corresponding to the maximum ΔK. Barplots were generated with DISTRUCT v1.1 (Rosenberg 2004).

## Results

### Mitochondrial 16S rDNA

Clear 16S rDNA sequences were obtained for 145 earthworms, 120 from the three sites and 25 from bait shops (Table 1). Sanger sequences of 15 individuals were of bad quality and were not included in the 16S rDNA dataset, but microsatellites amplified well for all individuals. Overall, the 16S rDNA sequences showed a maximum p-distance of 3.9%; and the 145 individual sequences were collapsed into 36 haplotypes (Table 1).

Three haplotypes were very common. The most common (18 individuals) was present in two bait shop individuals and in all subpopulations except MN_100_3. Two other haplotypes were also very common (both with 17 individuals), but were less widespread. One occurred in four subpopulations (MN_0_3, MN_50_1, MN_50_2, MN_100_1) and in one bait shop individual. The other was present in six subpopulations (MN_0_3, MN_50_1, MN_50_2, MN_50_3, MN_100_1, MN_100_3) and in one bait shop individual. Subpopulation MN_50_3 had the highest number of different haplotypes relative to the number of individuals sampled (14 individuals with nine haplotypes), followed by subpopulations MN_100_1 and MN_100_2 (seven haplotypes in twelve and 13 individuals, respectively). Subpopulation MN_100_3 had the lowest number of haplotypes, all but one individual had identical 16S rDNA sequences, the other haplotype in this subpopulation was genetically distinct (p-distance=1.9%) and co-occurred only in a single bait shop individual.

The haplotype network showed no clear structure among populations or bait-shop and field-collected individuals (Figure 2). Subpopulations included both unique and frequent haplotypes that cooccurred in other subpopulations. Earthworms obtained from bait shops carried six haplotypes that were not identical with field collected haplotypes, but also seven haplotypes that were identical with individuals from the field. The maximum likelihood analysis showed seven well-supported clades (bootstrap support >90%, Supplementary Information, Fig. S1), five of these clades included baitshop and field-collected individuals and two clades comprising five individuals that were exclusively sampled from bait shops but which were distinct from field-collected individuals.

**Figure 2.**
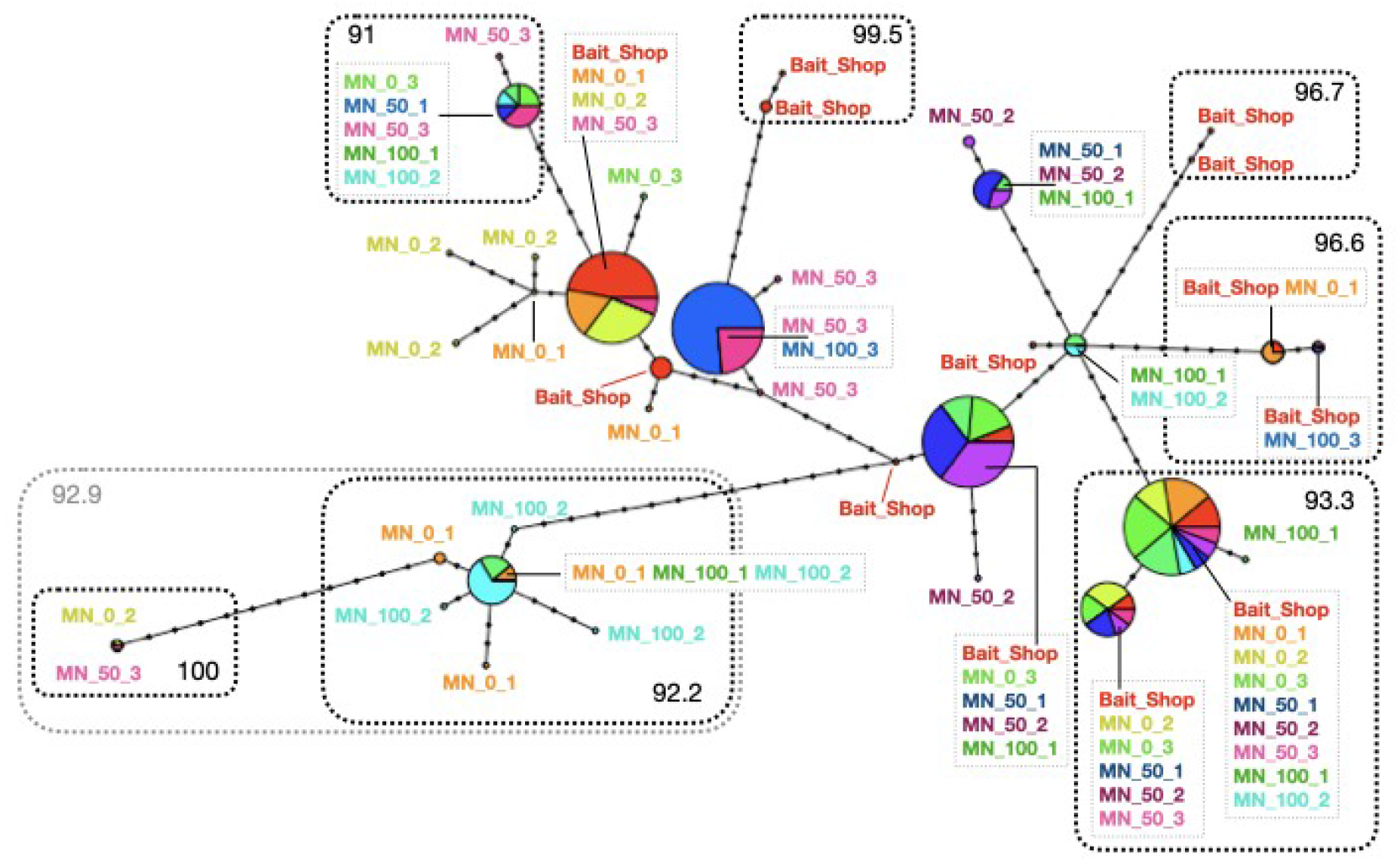
Haplotype network of 16S rDNA sequences collected from the field and bait shops in the St. Paul-Minneapolis area (MN). The network shows that seven distinct lineages are present in the field, some are rare (small circles representing one individual), others are widely distributed (large circles representing several individuals from different subpopulations). The genetic distinctness among haplotypes is reflected by the length of the lines connecting circles, and dots on lines represent hypothetical mutation steps. The size of circles is relative to the number of individuals represented by this haplotype. The dashed boxes outline well-supported clades from the Maximum Likelihood tree, black numbers in boxes are bootstrap support values.

### Nuclear microsatellites

The microsatellite dataset contained 160 individuals and eight loci. The number of alleles per locus across all populations was similar and ranged from 17 (LTM128) to 22 (LTM278), except for locus LTM252 which had 34 alleles (Supplementary Information, Fig. S2a). In all but two loci (LTM026 and LTM187), the observed heterozygosity was lower than the expected heterozygosity (Supplementary Information, Fig. S2b). Further, all loci deviated significantly from HWE and the mean observed heterozygosity varied significantly among loci (Bartlett’s test: p=<0.05).

Each field collected population consisted of 45 individuals (15 individuals per subpopulation, three subpopulations per site), and the pooled Bait_Shop population consisted of 25 individuals. Population MN_100 had the highest numbers of alleles (Na=117; Supplementary Information, Fig. S2c), closely followed by populations MN_0 (Na=112) and MN_50 (Na=111). Bait shop individuals had considerably fewer numbers of alleles (Na=90), but sample size was also lower. When comparing the ten subpopulations (Supplementary Information, Fig. S2d), MN_100_1 had the highest number of alleles (Na=87) and MN_100_3 had the lowest number of alleles (Na=62). The number of alleles of the three subpopulations at MN_0 and subpopulation MN_50_3 ranged between 72 and 75. The number of alleles of the remaining subpopulations were lower with Na=66 (MN_50_1) and Na=65 (MN_50_2 and MN_100_2). The global F_ST_ and F_IS_ values, that take all genotypes and all loci into account, were smaller for the four populations (Supplementary Information, Fig. S2e), with F_ST_=0.045 and F_IS_=0.089 compared to the ten subpopulations dataset (Supplementary Information, Fig. S2f) with values of F_ST_=0.068 and slightly lower F_IS_=0. 0.062.

The mean pairwise F_ST_ values between populations were also very close to zero, ranging between 0.033 and 0.058 (Supplementary Information, Table S1). However, the range of pairwise F_ST_ values was greater at subpopulation level, ranging between 0.013 and 0.139 (Table 2).

The AMOVA was significant at all levels and showed that genetic variance was quite low between populations and subpopulations. The majority of variance (89%) was found within samples (individuals). Variance between subpopulations within populations was low (4.9%), and variance between individuals within sites was even lower (3.7%). The variance was lowest between populations (2.0%).

The DAPC cross-validation peaked around 40 PC axes. Closer examination showed that the clustering patterns were consistent after reducing the number of PC axis around that value for both, four populations (33 PC axes retained, eigenvalues of LD1=181.3 and LD2=145.0) and for ten subpopulations (33 PC axes retained, eigenvalues of LD1=80.2 and LD2=74.3). Clustering patterns showed that most subpopulations were very similar and overlapped in one cluster (Fig. 3), except for subpopulation MN_100_3, which separated from all other subpopulations at the first axis, and bait shop individuals, which separated from all other subpopulations at the second axis. The analysis with four populations (Supplementary Information, Fig. S3) showed similar clustering patterns in which population MN_0 and MN_50 overlapped in one cluster, and MN_100 and bait shop individuals represented the most distinct genetic clusters on the first and second axes.

**Figure 3.**
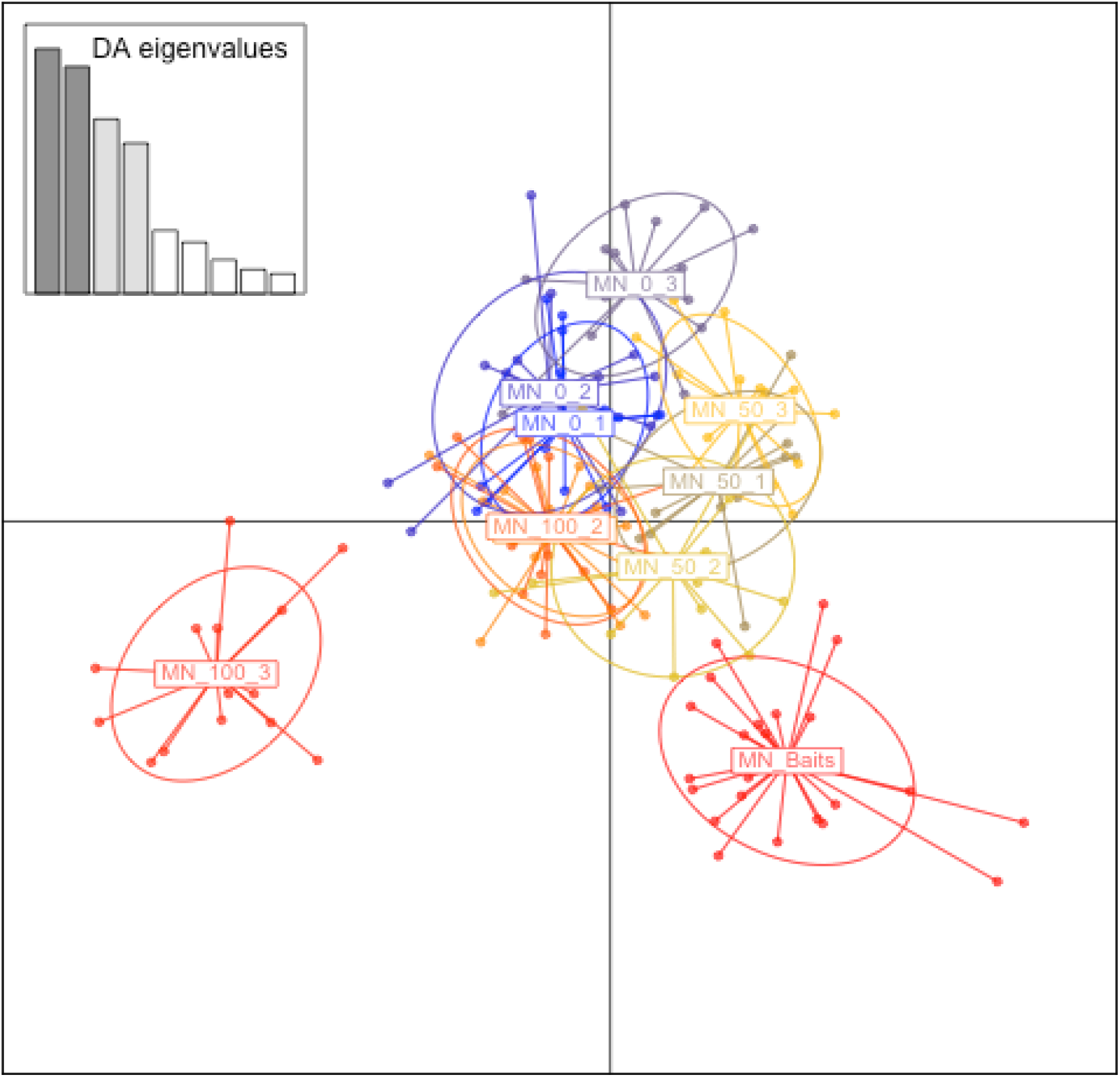
Discriminant analysis of principal components (DAPC) for 160 individuals of *Lumbricus terrestris* from ten subpopulations in the urban area of Minneapolis-St. Paul, MN (USA). The axes represent the first two Linear Discriminants (LD). Each circle represents a cluster and each dot represents an individual. The subpopulations weakly differentiated along the first two axes, but rather overlap with each other, except for subpopulation MN_100_3 and individuals collected from bait shops (MN_Baits).

Population assignment in STRUCTURE and STRUCTURE HARVESTER indicated that the sampling area most likely comprised five genetically differentiated populations (K=5, Fig. 4). Four clear patterns occurred. First, subpopulations 5 m apart (subpopulations _1 and _2 per sampling site) were most similar. Second, at sampling sites MN_50 and MN_100, subpopulations at 5000 m distance were considerably differentiated from the other two subpopulations at the same site. Third, sampling sites MN_0_1-3 and subpopulation MN_50_3 represented one genetic cluster, which was consistent across analyses with other K-values. Finally, subpopulation MN_100_3 and bait shop individuals (Bait_Shop) also represented discrete clusters.

**Figure 4.**
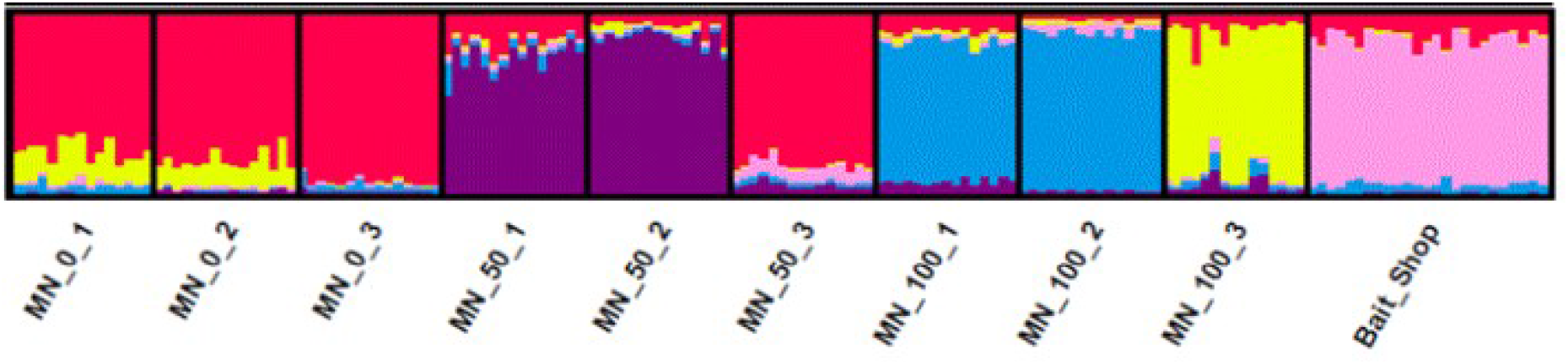
Bar plots depicting the assignment of individual genotypes of *L. terrestris* from ten subpopulations into five genetic clusters (K=5).

## Discussion

Using nuclear and mitochondrial markers, this study investigated the genetic structure of the invasive earthworm species *L. terrestris* in the metropolitan area of Minneapolis-St.Paul. Our results show that allelic differentiation and genetic variance of the nuclear markers was rather low between populations of *L. terrestris*, indicating that these populations are a homogenous group of genetically similar individuals. This suggests that these populations are established and likely interbreed. However, across all populations, the numbers of alleles and mitochondrial lineages was high, demonstrating that overall genetic diversity is substantial in *L. terrestris*, which is consistent with findings of earlier genetic studies in this species (Gailing et al. 2012; Klein et al. 2017; Keller et al. 2020). We found several mitochondrial lineages in each population, but genetic differentiation at nuclear markers was low, suggesting diffusive spread of earthworms across sampling sites followed by admixture. Diffusive spread describes a more continuous dispersal away from an initial source (Cameron et al. 2008). We could not identify hotspots of introductions, because several mitochondrial lineages were common and widespread among populations. This suggests that dispersal or short-distance transport of earthworms may occur at random in many directions, and that individuals may interbreed after transport with already present earthworms. Further, isolated mitochondrial haplotypes that were only present in single individuals were also common, suggesting recurrent transport or introductions of earthworms in the field which was not followed by successful breeding.

Alternative to multiple introductions with admixture, the high number of mitochondrial haplotypes accompanied by low genetic variance at nuclear markers within populations could be a result of the special reproductive mode of earthworms. *Lumbricus terrestris* is a hermaphrodite with reciprocal insemination, resulting in fertilisation of eggs and production of cocoons in each individual after mating (Edwards and Bohlen 1996). Consequently, it may be that a pair of earthworms may produce offspring that carries similar, admixed nuclear genomes but different mitochondrial lineages, due to the maternal transmission of mitochondria opposed to the biparental inheritance of nuclear genes. It is also possible that some earthworms invest more in sperm and less in eggs, or mate unsuccessfully, which may result in the presence of isolated mitochondrial lineages.

The population structure found for nuclear markers indicates that different dispersal processes acted on these earthworm populations. First, one subpopulation (MN_50_3) belonged genetically to a population 100 km south-east of its sampling site (MN_0), and not to the nearby subpopulations 5 km away, indicating effective jump dispersal, which is common in earthworms (Cameron et al. 2008, Klein et al. 2020). Second, one subpopulation in the north-west of the sampling area (MN_100_3) was also genetically differentiated from the nearby subpopulations 5 km away, but remained isolated in the population structure analyses and was dominated by one mitochondrial lineage, suggesting a recent founder event by passive dispersal of one or few individuals or a genetic bottleneck as this site is prone to recurrent flooding by the nearby St. Croix River. Third, individuals collected from bait shops separated genetically from all other subpopulations indicating that the free-living populations did not establish from recent or current cases of bait-dumping. Baits sold from shops that were not included in our sampling or from private vermicultures could also be a possible source for free-living earthworms. However, the high genetic similarity of earthworms sold in the five local bait shops suggests that local retailers purchased their products from a common source, such as a central distributor or earthworm farm. The genetic similarity of all bait shop individuals and their clear differentiation from free-living earthworms instead suggests that free-living earthworm populations derived from other sources or from releases of baits that were sold in the past. These patterns strongly contrast with findings from a study in Calgary, Alberta, Canada (Klein et al. 2017), which analysed *L. terrestris* populations from the field and bait shops using a similar study design and markers. They found that bait shop genotypes formed a genetic cluster with the urban population, indicating that bait disposal was an important source for free-living earthworm populations. Further, their populations were considerably more differentiated from each other than populations in this study, suggesting that free-living populations in Canada are rather recent introductions and with lower incidences of effective dispersal (dispersal with breeding), which is consistent with the more recent history of earthworm invasions in Alberta compared to Minnesota (Hale 2008, Klein et al. 2017).

The relevance of human activities for dispersal and establishment of earthworm populations is well known (Holdsworth et al. 2007; Keller et al. 2007; Cameron et al. 2007, 2008; Klein et al. 2017, 2020). The next step for investigating earthworm invasions is to disentangle the different anthropogenic vectors acting on local populations in order to manage these invasions. Once established, it is impossible to remove earthworm populations, and the development of management strategies to confine the invasion therefore remains the primary option when faced with established populations (Keller et al. 2007; Lawson Handley et al. 2011). The well-known link between baits and free-living populations (Holdsworth et al. 2007; Keller et al. 2007; Cameron et al. 2008; Klein et al. 2017) could not be identified in this study. Possibly, public awareness and educational campaigns reduced bait disposal in the past decade (Callaham et al. 2006; Keller et al. 2007). These campaigns are dedicated to education around earthworms and their effect and exist in Minnesota and the upper Midwest (e.g., https://wormwatch.d.umn.edu). They also distribute information to anglers about the importance of proper bait disposal. Other human-related factors that correlate with earthworm invasions are farming (Keller et al. 2007; Klein et al. 2020), logging and proximity to roads (Dymond et al 1997; Gundale et al. 2005; Suarez et al. 2006; Holdsworth et al. 2007; Hendrix et al. 2008; Cameron et al. 2008; Callaham et al. 2016; Roger and Collins 2017). Further, genetic differentiation between populations at small scale (0.6-13 km) is common in *L. terrestris* (Klein et al. 2017, Keller et al. 2020), but was not present in the population we sampled in the south from an area of intense agriculture (MN_0). Non-directional transport of earthworms with farm equipment and soil-movement by tillage practices could explain the genetic homogeneity and connectivity of this population, suggesting that farming facilitates the dispersal of *L. terrestris* and likely that of other invasive earthworm species such as *Aporrectodea caliginosa*, that are less sensitive to disturbances (Ivask et al. 2007; Smith et al. 2008).

Apart from identifying source populations and disentangling invasion routes, it is also important to understand what determines the success of invasive non-native species in order to implement successful management strategies (Lawson Handley et al. 2011). Here, earthworm populations consisted of both common and widespread, but also rare and isolated mitochondrial lineages, which is consistent with other molecular studies on invasive earthworm species in North America (Cameron et al. 2008; Gailing et al. 2012; Porco et al. 2013; Schult et al. 2016; Klein et al. 2017, 2020; Keller et al. 2017, 2020). Why some mitochondrial lineages thrive and reproduce and others do not depicts one of the most fundamental questions of invasion biology (Lawson Handley et al. 2011). Repeated sampling of populations that contained the most widespread haplotypes, but also of populations that contained many isolated haplotypes would allow to observe changes in the frequency of genetic lineages over time and thereby monitor the viability of earthworm lineages.

This study revealed that earthworm populations in the metropolitan region of Minneapolis-St. Paul, Minnesota, are established and that different patterns of population structure, introductions and dispersal co-occurred, demonstrating that molecular markers are a valuable tool to study invasion patterns of earthworms. However, this study also showed that additional sampling is required to disentangle different human-related mechanisms of earthworm dispersal. Transcriptome screening for selective sweeps in different genetic lineages could eventually help to identify whether different lineages acquired adaptations that correlate with local invasions and how populations cope with founder effects, bottlenecks and local inbreeding (Gailing et al. 2012, Klein et al. 2017, 2020, Keller et al. 2020). The advances in sequencing technologies provide a valuable toolkit to launch new studies on invasive earthworm populations in order to infer the invasion success of some lineages as opposed to others, and will ultimately aid in developing management strategies to limit earthworm invasions.

## Declarations

### Funding

This study was funded by the German Research Foundation (DFG grant: Ei 862/7-1, SCHA1671/5-1). NE acknowledges funding by the European Research Council (European Union’s Horizon 2020 research and innovation program; grant no 677232) and the German Centre for Integrative Biodiversity Research Halle-Jena-Leipzig, funded by the German Research Foundation (DFG; FZT 118, 202548816).

### Conflicts of interest/Competing interests

No conflicts of interest are declared.

### Availability of data and material

All 16S rDNA sequences are available at GenBank (www.ncbi.nlm.nih.gov) under the accession numbers ON506389 to ON506413 (bait shop individuals) and ON506269 to ON506388 (field collected animals). Please, also check information in Code availability.

### Code availability

The R code used for data analysis is available at Dryad (doi:10.5061/dryad.w0vt4b8v2) or from the corresponding author on request. The files include the following data and analyses: aligned 16S rDNA dataset of this study for phylogeny and haplotype networks, microsatellite data and analyses of this study and microsatellite data from Klein et al. 2017 to compare population structures.

### Ethics approval

Not applicable

### Authors’ contribution

IS, SS, NE conceived and designed the study, AR, AK and BH did the field work, BH performed the laboratory work and analysed the data. BH and IS drafted the manuscript. All authors read, edited, and approved the final manuscript.

## Acknowledgments

We are grateful to Sabrina Pach for her support in genotyping the microsatellites at the Veterinary Medicine (University of Göttingen, Germany) and for permissions to collect specimens which were provided by the Department of Natural Resources Minnesota and the Warner Nature Center.

## Notes

### Competing Interest Statement

The authors have declared no competing interest.

## References

Addison JA (2009) Distribution and impacts of invasive earthworms in Canadian forest ecosystems. Biol Inv 11:59–79.

Arcese P, Rodewald AD (2019) Predictors and consequences of earthworm invasion in a coastal archipelago. Biol Inv 21:1833–1842.

Bienert F, De Danieli S, Michel C, Coissac E, Poillot C, Brun J-J, Taberlet P (2012) Tracking earthworm communities from soil DNA. Mol Ecol 21:2017–2030.

Callaham MA, González G, Hale CM, Heneghan L, Lachnicht SL, Zou X (2006) Policy and management responses to earthworm invasions in North America. Biol Inv 8:13317–1329.

Callaham MA, Snyder BA, James SW, Oberg ET (2016) Evidence for ongoing introduction of non-native earthworms in the Washington, DC metropolitan area. Biol Inv 18:3133–3136.

Cameron EK, Bayne EM, Clapperton MJ (2007) Human-facilitated invasion of exotic earthworms into northern boreal forests. Ecoscience 14:482–490.

Cameron EK, Bayne EM, Coltman DW (2008) Genetic structure of invasive earthworms *Dendrobaena octaedra* in the boreal forest of Alberta: insights into introduction mechanisms. Mol Ecol 17:1189–1197.

Craven D, Thakur MP, Cameron E et al (2017) The unseen invaders: introduced earthworms as drivers of change in plant communities in North American forests (a meta-analysis). Global Change Biol 23:1065–1074

Crumsey JM, Capowiez Y, Goodsitt MM, Larson S, Le Moine JM, Bird JA, Kling GW, Nadelhoffer KJ (2015) Exotic earthworm community composition interacts with soil texture to affect redistribution and retention of litter-derived C and N in northern temperate forest soils. Biogeochemistry 126:379–395.

Dobson AM, Blossey B, Richardson JB (2017) Invasive earthworms change nutrient availability and uptake by forest understory plants. Plant Soil 421:175–190.

Drouin M, Bradley R, Lapointe L, Whalen J (2014) Non-native anecic earthworms (*Lumbricus terrestris* L.) reduce seed germination and seedling survival of temperate and boreal tree species. Appl Soil Ecol 75:145–149.

Dymond P, Scheu S, Parkinson D (1997) Density and distribution of *Dendrobaena octaedra* (Lumbricidae) in aspen and pine forests in the Canadian Rocky Mountains (Alberta). Soil Biol Biochem 29:265–273.

Eisenhauer N, Partsch S, Parkinson D, Scheu S (2007) Soil Biol Biochem 39:1099–1110.

Edwards CA, Bohlen PJ (1996) Biology and ecology of earthworms. Chapman and Hall, London.

Evanno, G., Regnaut, s., Goudet, J. (2005) Detecting the number of clusters of individuals using the software STRUCTURE: a simulation study. Mol Ecol 14:2611–2620.

Ferlian O, Eisenhauer N, Aguirrebengoa M, Camara M, Ramirez-Rojas I, Santos F, Tanalgo K, Thakur MP (2018) Invasive earthworms erode soil biodiversity: A meta-analysis. J Animal Ecol 87:162–172.

Ferlian O, Thakur MP, Castañeda González A, San Emeterio LM, Marr S, Da Silva Rocha B, Eisenhauer N (2020) Soil chemistry turned upside down: a meta-analysis of invasive earthworm effects on soil chemical properties. Ecology 101:e02936.

Fernandez R, Novo M, Marchán DF, Díaz Cosín DJ (2016) Diversification patterns in cosmopolitan earthworms: similar mode but different tempo. Mol Phyl Evol 94:701–708.

Fisichelli NA, Frelich LE, Reich PB, Eisenhauer N (2013) Linking direct and indirect pathways mediating earthworms, deer, and understory composition in Great Lakes forests. Biol Inv 15:1057–1066.

Fisichelli NA, Miller KM (2018) Weeds, worms, and deer: positive relationships among common forest understory stressors. Biol Inv 20:1337–1348.

Frelich LE, Blossey B, Cameron EK, Dávalos A, Eisenhauer N, Fahey T, Ferlian O, Groffmann PM, Larson E, Loss SR, Maerz JC, Nuzzo V, Yoo K, Reiche PB (2019) Side-swiped: Ecological cascades emanating from earthworm invasion. Front Ecol Environ 17:502–510.

Gailing O, Hickey E, Lilleskov E, Szlavecz K, Richter K, Potthoff M (2012) Genetic comparisons between North American and European populations of *Lumbricus terrestris* L. Biochem Syst Ecol 45:23–30.

Gates GE (1978) The earthworm genus Lumbricus in North America. Megadrilogica 3:81–116.

Goudet J, Jombart T (2020) hierfstat: estimation and tests of hierarchical F-statistics. R package version 0.5-7.

Gundale MJ, Jolly WM, Deluca TH (2005) Susceptibility of a northern hardwood forest to exotic earthworm invasion. Cons Biol 19:1075–1083.

Gunn A. (1992) The use of mustard to estimate earthworm populations. Pedobiologia 36:65–67.

Hale CM (2008) Evidence for human-mediated dispersal of exotic earthworms: support for exploring strategies to limit further spread. Mol Ecol 17:1165–1169.

Hale CM, Frelich LE, Reich PB (2005) Exotic European earthworm invasion dynamics in northern hardwood forests of Minnesota, USA. Ecol Appl 15:848–860.

Hendrix PF, Baker GH, Callaham Jr MA, Damoff GA, Fragoso C, González G, James SW, Lachnicht SL, Winsome T, Zou W (2006) Invasion of exotic earthworms into ecosystems inhabited by native earthworms. Biol Inv 8:1287–1300.

Hendrix PF, Callaham Jr MA, Drake JM, Huang C-Y, James SW, Snyder BA, Zhang W (2008) Pandora’s box contained bait: the global problem of introduced earthworms. Annu Rev Ecol Evol Syst 39:593–613.

Holdsworth AR, Relich LE, Reich PB (2007) Effects of earthworm invasion on plant species richness in Northern hardwood forests. Cons Biol 21:997–1008.

Ivask M, Kuu A, Sizov E (2007) Abundance of earthworm species in Estonian arable soils. Europ J Soil Biol S39–S42.

Jakobsson M, Rosenberg NA (2007) CLUMM: a cluster matching and permutation program for dealing with label switching and multimodality in analysis of population structure. Bioinformatics 23:1801–1806.

Jombart T (2008) adegenet: a R package for the multivariate analysis of genetic markers. Bioinformatics 24:1403–1405.

Jombart T, Devillard S and Balloux F (2010) Discriminant analysis of principal components: a new method for the analysis of genetically structured populations. BMC Genetics 11:94.

Jombart T, Ahmed I (2011) adegenet 1.3-1: new tools for the analysis of genome-wide SNP data. Bioinformatics 27:3070–3071.

Kamvar ZN, Tabima FJ, Grünwald NJ (2014) poppr: an R package for genetic analysis of populations with clonal, partially clonal, and/or sexual reproduction. PeerJ 2:e281.

Kamvar ZN, Brooks JC, Grünwald NJ (2015) Novel R tools for analysis of genome-wide population genetic data with emphasis on clonality. Front Genet 6:208.

Keller RP, Cox AN, van Loon C, Lodge DM, Herborg L-M, Rothlisberger J (2007) From bait shops to the forest floor: earthworm use and disposal by anglers. Am Midl Nat 158:321–328.

Keller EL, Göres JH, Schall JJ (2017) Genetic structure of wo invasive earthworms, *Amynthas agrestis* and *Amynthas tokioensis* (Oligochaeta, Megascolecidae), and a molecular method for species identification. Megadrilogica 22:140–149.

Keller EL, Conolly ST, Görres JH, Schall JJ (2020) Genetic diversity of an invasive earthworm, *Lumbricus terrestris*, at a long-term trading crossroad, the Champlain Valley of Vermont, USA. Biol Inv 22:1723–1735.

Klein A, Cameron EK, Heimburger B, Eisenhauer N, Scheu S, Schaefer I (2017) Changes in the genetic structure of an invasive earthworm species (*Lumbricus terrestris*, Lumbricidae) along an urban-rural gradient in North America. Appl Soil Ecol 120:265–272.

Klein A, Eisenhauer N, Schaefer I (2020) Invasive lumbricid earthworms in North America – different life histories but common dispersal? J Biogeogr 47:674–685.

Larsson A (2014) AliView: a fast and lightweight alignment viewer and editor for large datasets. Bioinformatics 30:3276–3278.

Lawson Handley Estoup A, Evans DM, Thomas CE, Lombaert E, Facon B, Aebi A, Roy HE (2011) Ecological genetics of invasive alien species. Biol Control 56:409–428.

Nuzzo VA, Maerz JC, Blossey B (2009) Earthworm invasion as the driving force behind plant invasion and community change in Northeastern North American forests. Cons Biol 23:966–974.

Nuzzo V, Dávalos A, Blossey B (2015) Invasive earthworms shape forest seed bank composition. Diversity Distrib 21:560–570.

Paradis E (2010) pegas: an R package for population genetics with an integrated-modular approach. Bioinformatics 26:419–420.

Paradis E, Schliep K (2019) ape 5.0: an environment for modern phylogenetics and evolutionary analyses in R. Bioinformatics 35:526–528.

Paudel S, Longcore T, MacDonal B, McCormick MK, Szlavecz K, Wilson GWT, Loss SR (2016) Belowground interactions with aboveground consequences: invasive earthworms and arbuscular mycorrhizal fungi. Ecology 97:605–614.

Pérez-Losada M, Ricoyb M, Marscshal JC, Domíngues J (2009) Phylogenetic assessment of the earthworm *Aporrectodea caliginosa* species complex (Oligochaeta: Lumbricidae) based on mitochondrial and nuclear DNA sequences. Mol Phyl Evol 52:293–302.

Porco D, Decaens T, Deharveng L, James SW, Skarzynski D, Erséus C, Butt KR, Richard B, Hebert PDN (2013) Biological invasions in soil: DNA barcoding as a monitoring tool in a multiple taxa survey targeting European earthworms and springtails in North America. Biol Inv 15:899–910.

Potvin LR, Lilleskov EA (2017) Introduced earthworm species exhibited unique patterns of seasonal activity and vertical distribution, and *Lumbricus terrestris* burrows remained usable for at least 7 years in hardwood and pine stands. Biol Fertil Soils 53:187–198.

Pritchard JK, Stephens M, Donnelly P (2000) Inference of population structure using genotype data. Genetics 155:945–959.

Resner K, Yoo K, Sebestyen SD, Aufdenkampe A, Hale C, Lyttle A, Blum A (2015) Invasive earthworms deplete key soil inorganic nutrients (Ca, Mg, K, and P) in a Northern hardwood forest. Ecosystems 18:89–102.

Reynolds JW, Linden DR, Hale CM (2002) The earthworms of Minnesota (Oligochaeta: Acanthrodrilidae: Lumbricidae and Megascolecidae). Megadrilogica 8:85–100.

Rogers JA, Collins CD (2017) Ecological predictors and consequences of non-native earthworms in Kennebec County, Maine. Northeastern Naturalist 24:121–136.

Rosenberg NA (2004) DISTRUCT: a program for the graphical display of population structure. Mol Ecol Notes 4:137–138.

Schliep KP (2011) phangorn: phylogenetic analysis in R. Bioinformatics 27:592–593.

Schult N, Pittenger K, Davalos S, McHugh D (2016) Phylogeographic analysis of invasive Asian earthworms (*Amynthas*) in the northeast United States. Invertebrate Biol 135:314–327.

Sims RW, Gerard BM (1999) Earthworms-Synopses of the British Fauna, No 31. Dorchester, Great Britain: The Dorset Press.

Smith RG, McSwiney CP, Grandy AS, Suwanawaree P, Snider RM, Robertson GP (2008) Diversity and abundance of earthworms across an agricultural land-use intensity gradient. Soil Tillage Res 100:83–88.

Suárez ER, Tierney GL, Fahey TJ, Fahey R (2006) Exploring patterns of exotic earthworm distribution in a temperate hardwood forest in south-central New York, USA. Landscape Ecol 21:297–306.

Tiunov AV, Hale CM, Holdsworth AR, Vsevolodova-Perel TS (2006) Invasion patterns of Lumbricidae into the previously earthworm-free areas of northeastern Europe and the western Great Lakes region of North America. Biol Inv 8:1223–1234.

van Oosterhaut C, Hutchinson WF, Willis DPM, Shipley P (2004) MICRO-CHECKER: software for identifying and correcting genotyping errors in microsatellite data. Mol Ecol Notes 4:535–538.

Velavan TP, Schulenburg H, Michiels NK (2007) Development and characterization of novel microsatellite markers for the common earthworm (*Lumbricus terrestris* L.). Mol Ecol Notes 7:1060–1062.

Villesen P (2007) FaBox: an online toolbox for fasta sequences. Mol Ecol Notes 7:965–968.

