## Supplementary Information for "Population structure and genetic variance among local populations of an non-native earthworm species in Minnesota, USA"

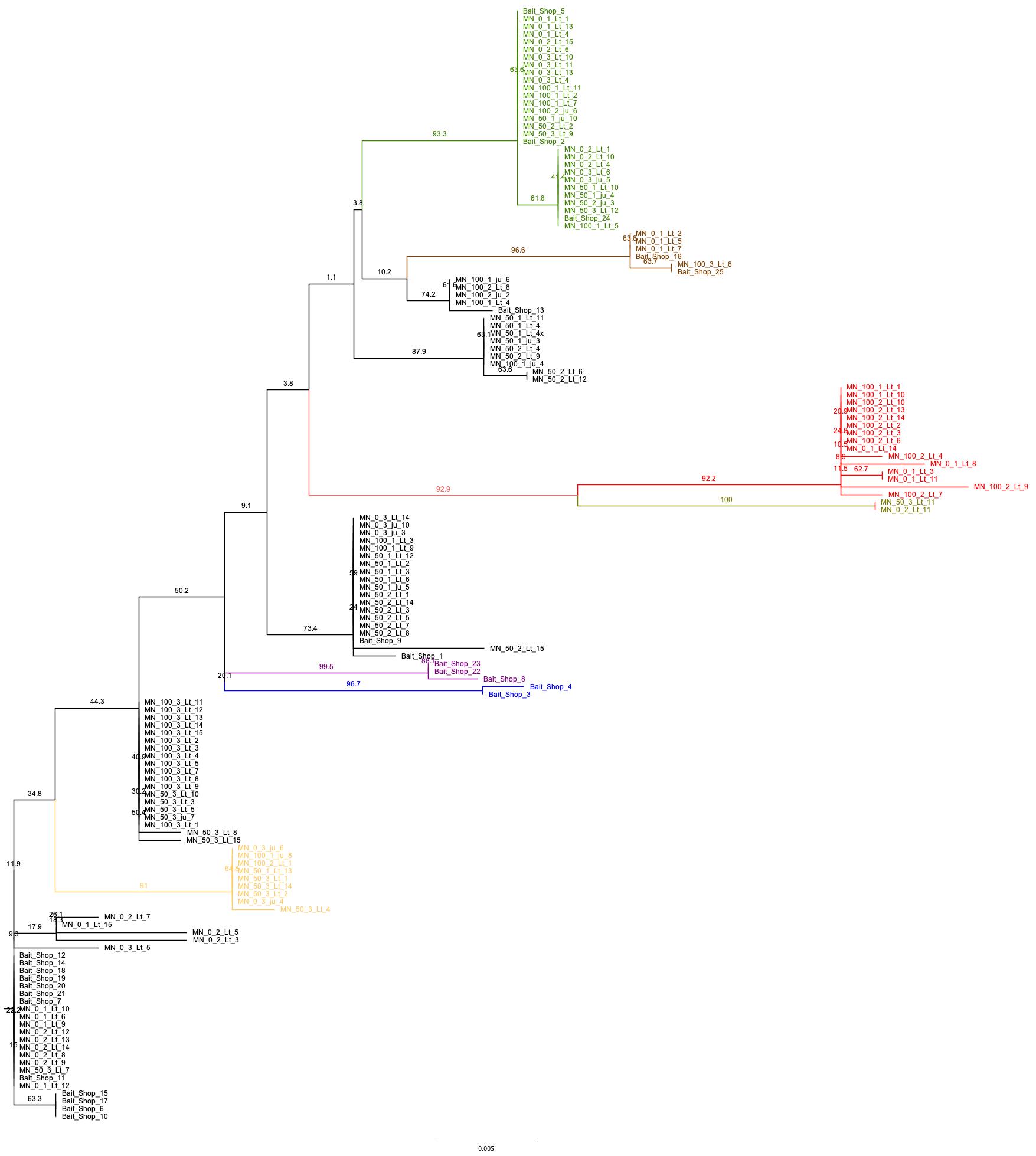

**Figure S1.** Maximum Likelihood tree of *Lumbricus terrestris* from three sites in the metropolitan area of Minneapolis-St. Paul (Minnesota, USA) based on 16S rDNA sequences, calculated with ape and phangorn packages in R. Node support is based on 1,000 bootstrap replicates. Well-supported clades are highlighted in different colours and correspond with boxes in the haplotype network of Fig. 2.

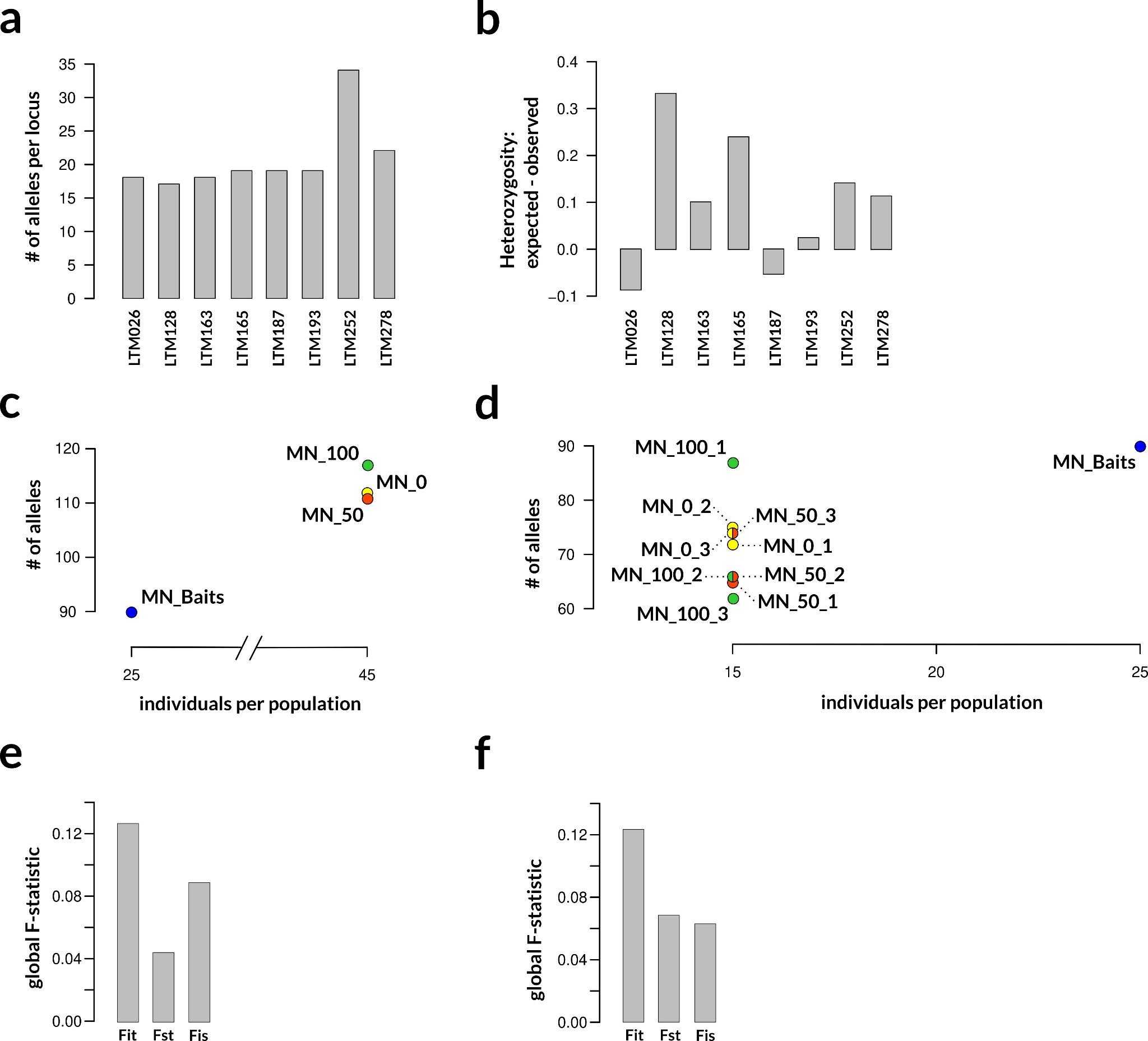

**Figure S2.** Summary of the microsatellite dataset showing (a) the number of alleles per locus, (b) the degree of heterozygosity per locus, the number of alleles for (c) four populations and (d) ten subpopulations, the global F-statistics for (e) four populations and (f) ten subpopulations.

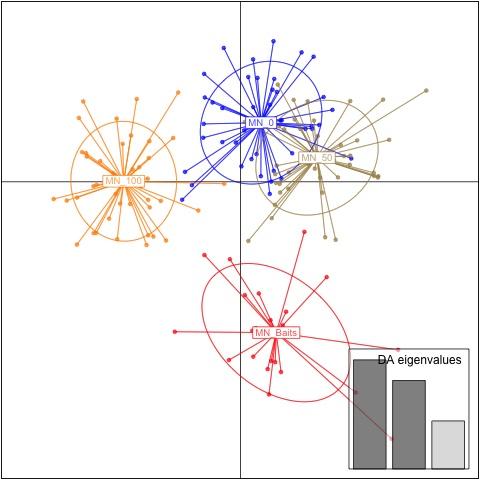

**Figure S3.** Discriminant analysis of principal components (DAPC) for 160 individuals of *L. terrestris* from four populations in the urban area of Minneapolis-St. Paul, MN (USA). The axes represent the first two Linear Discriminants (LD). Each circle represents a cluster and each dot represents an individual. Populations MN_0 and MN_50 largely overlap, supporting the genetic similarity of individuals in these populations. Population MN_100 separates on the first axis and MN_Baits on the second axis, which is in accordance of the higher Fst values of these populations to MN_0 and MN_40 (Table S1).

**Table S1.** Mean pairwise F_ST_ of *L. terrestris* between three field collected populations and bait shop individuals.

|  | **MN_0** | **MN_50** | **MN_100** |
| --- | --- | --- | --- |
| **MN_50** | 0.033 |  |  |
| **MN_100** | 0.049 | 0.044 |  |
| **Bait Shops** | 0.058 | 0.046 | 0.05 |
